# Identification of transposable element families from pangenome polymorphisms

**DOI:** 10.1101/2024.04.05.588311

**Authors:** Pío Sierra, Richard Durbin

## Abstract

**Background:** Transposable Elements (TEs) are fragments of DNA, typically a few hundred base pairs up to several tens of thousands bases long, that have the ability to generate new copies of themselves in the genome. Most existing methods used to identify TEs in a newly sequenced genome are based on their repetitive character, together with detection based on homology and structural features. As new high quality assemblies become more common, including the availability of multiple independent assemblies from the same species, an alternative strategy for identification of TE families becomes possible in which we focus on the polymorphism at insertion sites caused by TE mobility.

**Results:** We develop the idea of using the structural polymorphisms found in pangenomes to create a library of the TE families recently active in a species, or in a closely related group of species. We present a tool, **pantera**, that achieves this task, and illustrate its use both on species with well-curated libraries, and on new assemblies.

**Conclusions:** Our results show that **pantera** is sensitive and accurate, tending to correctly identify complete elements with precise boundaries, and is particularly well suited to detect larger, low copy number TEs that are often undetected with existing de novo methods.

## Background

Transposable Elements (TEs) often take up a large fraction of a eukaryotic genome, and they can also vary enormously between species (Wells and Feschotte, 2020). Due to their repetitive nature, TEs represent a problem for accurate genome assembly and read mapping, in particular when dealing with short reads. For that reason they have frequently been disregarded in many analyses, typically by masking them out (Smit, Hubley, R & Green, P., 2013). However, with the new longer read assembly protocols (Li and Durbin, 2024) it is now much easier to sequence through them so they are correctly represented in genome assemblies, making it easier to identify and classify them, and consequently to study them for their own sake.

The construction of a library of transposable elements present in an organism typically involves the following steps: search for elements homologous to those of an existing library for a closely related species; search for repetitive elements in a genome; search within these repetitive elements for defining structural features, such as LTRs (Long Terminal Repeats), TIRs (Terminal Inverted Repeats), known motifs and ORFs (Open Reading Frames) for characteristic proteins (Storer et al., 2022). TE Hub (Elliott et al., 2021) currently lists 51 tools associated with “library generation”. Two of the most popular ones are RepeatModeler (Flynn et al., 2020) and REPET (Flutre et al., 2011). There are also composite methods that merge the results of multiple other tools in pipelines, such as EarlGrey (Baril et al., 2022), PiRATE (Berthelier et al., 2018), TransposonUltimate (Riehl et al., 2022) and MCHelper (Orozco-Arias et al., 2023). It is also possible to process several genomes serially and combine their annotations to produce a more complete library (Ou et al., 2022). However essentially all these tools carry out their search starting from the set of chromosomal or contig sequences of a single assembled genome, or the raw sequencing reads for a single genome (Novák et al., 2010). In general these methods are characterised by a “repeat first” approach, in that they first look for dispersed repetitive sequences or structural motifs and then try to cluster them and extend them to recreate finally the complete sequence of the original TE. Here we instead take a “polymorphism first” approach, focusing on insertion polymorphisms between different copies of the genome as likely candidates for full length TEs, as described below.

Regardless of the method employed, to obtain a high quality library usually requires manual curation of the results, to ensure that the consensus sequences belong to TEs and not to other repeat types such as simple repeats or repeated gene families, and also to confirm that they represent the complete element, rather than just a fragment of it or an extension of it containing adjacent non-TE sequence (Goubert et al., 2022). Naturally, the extent to which the initial detection tool itself automatically creates full length consensus sequences for candidate transposon families is very important in determining the amount of effort required to curate the library.

One of the changes that makes it possible to look with a new light at TE biology is the existence of large scale projects like the Darwin Tree of Life (The Darwin Tree of Life Project Consortium, 2022) or Zoonomia (Genereux et al., 2020), which are rapidly increasing the number of species for which one or more high quality assemblies are available. The latest assemblers, like HiFiAsm (Cheng et al., 2021) or VERKKO (Rautiainen et al., 2023), aim to independently assemble the two haplotypes of a diploid genome. This provides multiple copies of genome sequences from the species that can be compared to identify structural polymorphisms at several levels. First, we can compare the two haplotypes of the same sample (or more if it is polyploid) to obtain heterozygous polymorphisms. Second, we can compare the genomes from two or more individuals of the same species to obtain intraspecies polymorphisms. Lastly, we can compare the genomes of two or more closely related species with a high level of synteny to obtain interspecies polymorphisms.

The other important change is the appearance of faster and more accurate whole genome alignment tools (Li, 2018) (Marco-Sola et al., 2022). In particular methods have been developed to generate pangenomes which represent multiple closely related genome sequences in a computational graph structure which explicitly identifies sequence segments present only in a subset of the genomes, i.e. structural variants (SVs) (Hickey et al., 2020) (Li et al., 2020) (Garrison et al., 2023). Projects such as the Human Pangenome Reference Consortium (Liao et al., 2023) are generating large pangenome graphs of this nature. A new tool, GraffiTE (Groza et al., 2023), has been developed to use these pangenomes to genotype transposable element insertion polymorphisms, starting from an existing TE library for the species under consideration. In our work we follow the opposite direction to show how a pangenome can be used to obtain a new TE library whose elements will be close in most cases to the complete sequence of the TE, even before any manual curation. Recently a related method has been made available, which starts from short read whole genome shotgun data to identify polymorphic structural variants as candidates for TEs (Burke et al., 2023).

## Methods

Our hypothesis is that most of the SVs in the size range of hundreds to tens of thousands of bases that we detect when comparing two or more closely related genomes are caused by TE insertions or deletions, as illustrated in Figure 1a,b. We identify candidate TE families by clustering the inserted sequences with high stringency for both sequence identity (default 95%) and full length alignment. Our tool returns consensus sequences of these putative TE families. These can later be classified into any of the more than 500 current superfamilies described in the literature using an existing classifier, in our case RepeatClassifier (Flynn et al., 2020). They can also be used directly with tools to annotate or mask TE copies in a genome, such as RepeatMasker (Flynn et al., 2020). Figure 1c provides an overview of the process, with Figure 1d illustrating the resulting annotation created by RepeatMasker for the *Drosophila melanogaster* genome, and Figure 1e,f,g an example of a specific identified element. Source code is available at github.com/piosierra/pantera.

**Figure 1.**
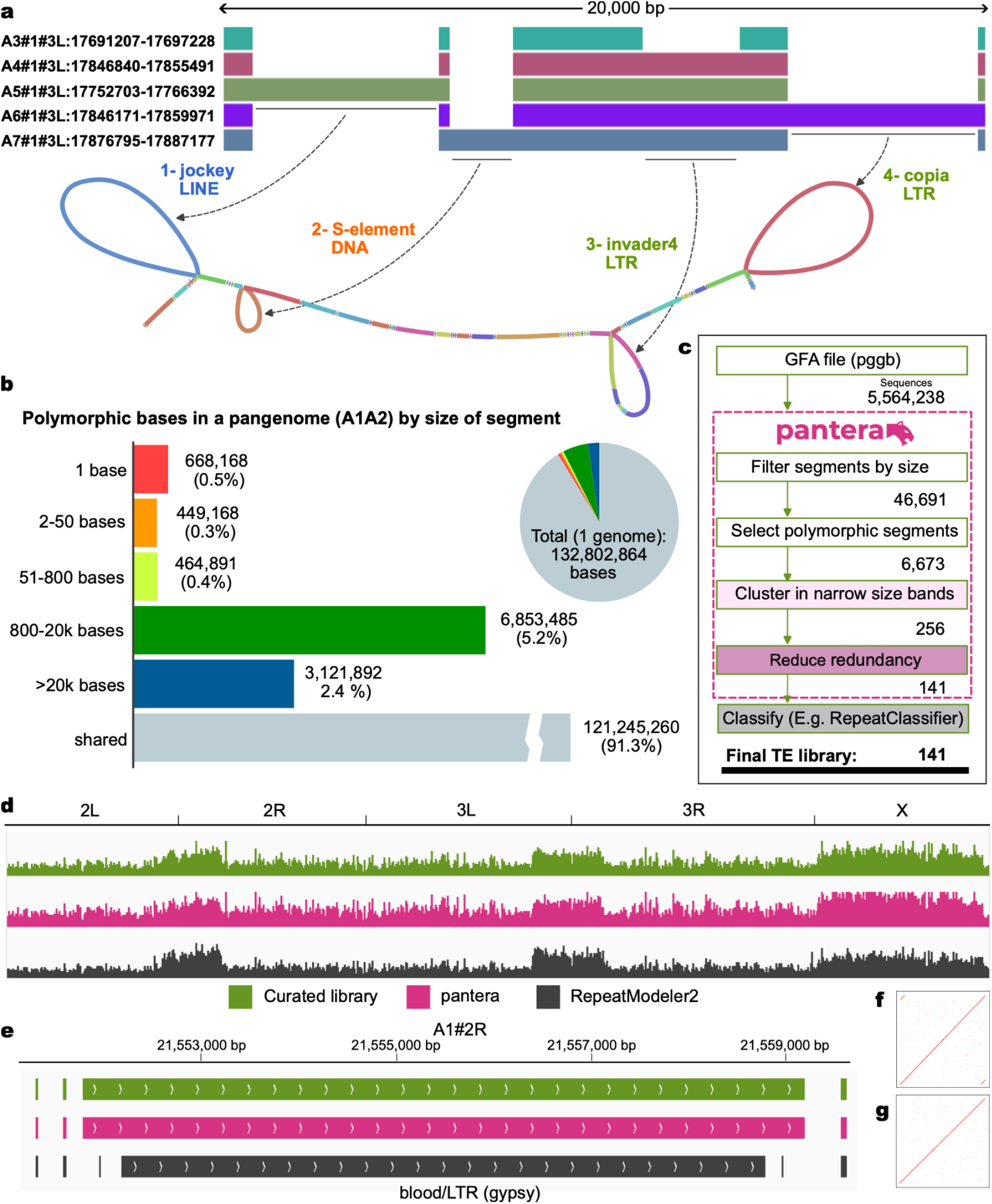
Obtaining a transposable element library from a pangenome. **a**, A 20kb section of a pangenome of chromosome 2R from seven high quality genomes of *Drosophila melanogaster* (just five shown, figure generated with odgi). Gaps in the bands represent structural variants, i.e. insertions or deletions in some of the genomes compared to the others. These structural variants can also be visualised as loops or “bubbles” in the graph representation. Here we see four structural variants each thousands of bases long, arising from four different TE insertions. **b**, Number of bases in a pangenome of two *Drosophila melanogaster* genomes (A1 and A2) by whether the bases are fully aligned (shared) or they do not align, binned by the size of the insertion or mismatch. **c**, Workflow of **pantera.** First it selects from the gfa file segments that are polymorphic and may hence belong to a TE. To reduce the number of false positives only segments for which there are at least two almost identical polymorphic sequences are selected (cluster in narrow size bands). Then, a less stringent clustering is performed to reduce redundancy and generate the final TE library that can be classified with any existing tools. **d**, Annotations of the A1 *Drosophila melanogaster* genome obtained with RepeatMasker using three different libraries. Green: curated reference library. Pink: pantera *de novo* library. Grey: RepeatModeler *de novo* library. **e**, Example of an LTR element (Blood) for which pantera was able to correctly identify the full element, including its LTR components (**f**) that in this case are not fully reported by RepeatModeler, neither as part of the full consensus (**g**) nor as a solo LTR element.

### Prerequisites

The only required input to run pantera is a pangenome in the form of one or several GFA files, that include the individual paths on the graph as P entries or the links as L entries, all with non overlapping links between segments. We have generated these files with pggb (Garrison et al., 2023), but other pangenome graph generators should also be usable. For the pangenomes described in this paper we used pggb parameters -p 85 to 95 (minimum average nucleotide identity for a seed mapping) and -s 2000 to 5000 (segment length), depending on the expected levels of identity of the sequences included, and workflow versions wfmash v0.9.1-3-gc5882a1 “Mutamento”, seqwish v0.7.6 “Temporaneo”, odgi v0.7.3 “Fissaggio”. No master reference is needed. When the pangenome is divided into several GFA files, for example one for each chromosome, pantera can take a list of GFA files to process.

### Segment selection and clustering

The first step is to select just the segments in the graph that belong to polymorphisms. In the case of a graph made from just two genomes, such as the haplotypes of a diploid genome, we directly select segments that belong to just one path and are flanked by segments shared by both paths. To avoid potential artefacts on the pangenome we select only those for which the flanking shared segments are larger than a certain size (default = 1). Furthermore, we require that those flanking segments are linked together in the alternative path. This prevents the selection of segments which do not belong to a polymorphic insertion, but rather to a very divergent section. When the graph contains more than two paths, we apply this rule to each pair of paths. Next we filter these segments to select only those falling in a certain length range (defaults 250 to 50,000 bases).

Once the segments are filtered they are ordered by length and allocated into bins based on overlapping length windows of width 100 bp. These bins are then clustered using an R implementation of the cd-hit algorithm (McDavid et al., 2023), with default parameters of 300 maximum number of sequences to cluster and 95% of identity (cd-hit-est options: -c 0.95 -G 1 -g 1). Sequences from clusters with at least a given number of sequences (default = 2) are then selected and any of the redundancy from the overlapping windows removed. Then they are clustered again, with wider windows of width 2,000 bp, and the resulting clusters are aligned with mafft. A consensus is then obtained from the aligned sequences, using a stringent plurality threshold of 60% on the first pass that helps remove random edge sequences.

After producing the initial library we remove subfragments that are contained within complete TEs, such as solo LTRs or partial LINE elements. This is done by repeating the clustering process, reducing the identity needed to cluster (default on second pass 85%) but increasing the requirements of coverage for the smaller sequence aligned (cd-hit-est options: -c 0.85 -G 0 -aS 0.90 -uL 0.05 -g 1).

### TE family classification

The previous steps produce a set of sequences which are expected to belong to transposable element families. They can then be classified using any TE classifier tool, for example RepeatClassifier (Flynn et al., 2020). We also used CENSOR (Kohany et al., 2006) with some of the families to examine their homology to previously identified sequences.

To assess the presence of standard TE features, open reading frames (ORFs) for the sequences were obtained with getorf (Rice et al., 2000) and structural features (LTRs, TIRs, polyA tail) detected with our own script pantercheck, in which TE candidate sequences are blasted to themselves one by one, allowing one mismatch for each 8 bases. Elements are defined as “complete” as follows: 1) for DNA elements (except Cryptons) and LTR DIRS elements, having terminal inverted repeat (TIR) sequences at least 10 bases long;. 2) for LTR elements, having long terminal repeat (LTR) sequences at least 10 bases long; 3) for LINE elements, having a candidate ORF at least 1,300 aminoacids long, and a polyA tail. In addition LINE1 elements required a further ORF1 candidate at least 700 aminoacids long. As RepeatModeler returns results for LTR elements as two components, internal and LTR, when scoring RepeatModeler results we count as “complete” any LTR result with an internal segment larger than 3,000 bases that also has the corresponding LTR segment. The remainder of the LTR results are considered as incomplete. These can include solo LTR segments for which a copy of the full LTR element is no longer present in the genome.

## Results

First, we benchmarked the new tool with three species for which there exist manually curated TE libraries: *Drosophila melanogaster* (fruit fly)*, Oryza sativa* (rice), *Danio rerio* (zebrafish). Figure 2 shows the distribution of insertion polymorphisms in the size range 250bp to 20kb in the pangenomes we used for these species, with the fraction of those polymorphisms exhibiting the various TE features classified by pantercheck. For each of the species we compared the results of pantera to those of a reference library and those obtained with a denovo method, RepeatModeler for *Drosophila melanogaster* and *Danio rerio* and REPET for *Oryza sativa*.

**Figure 2:**
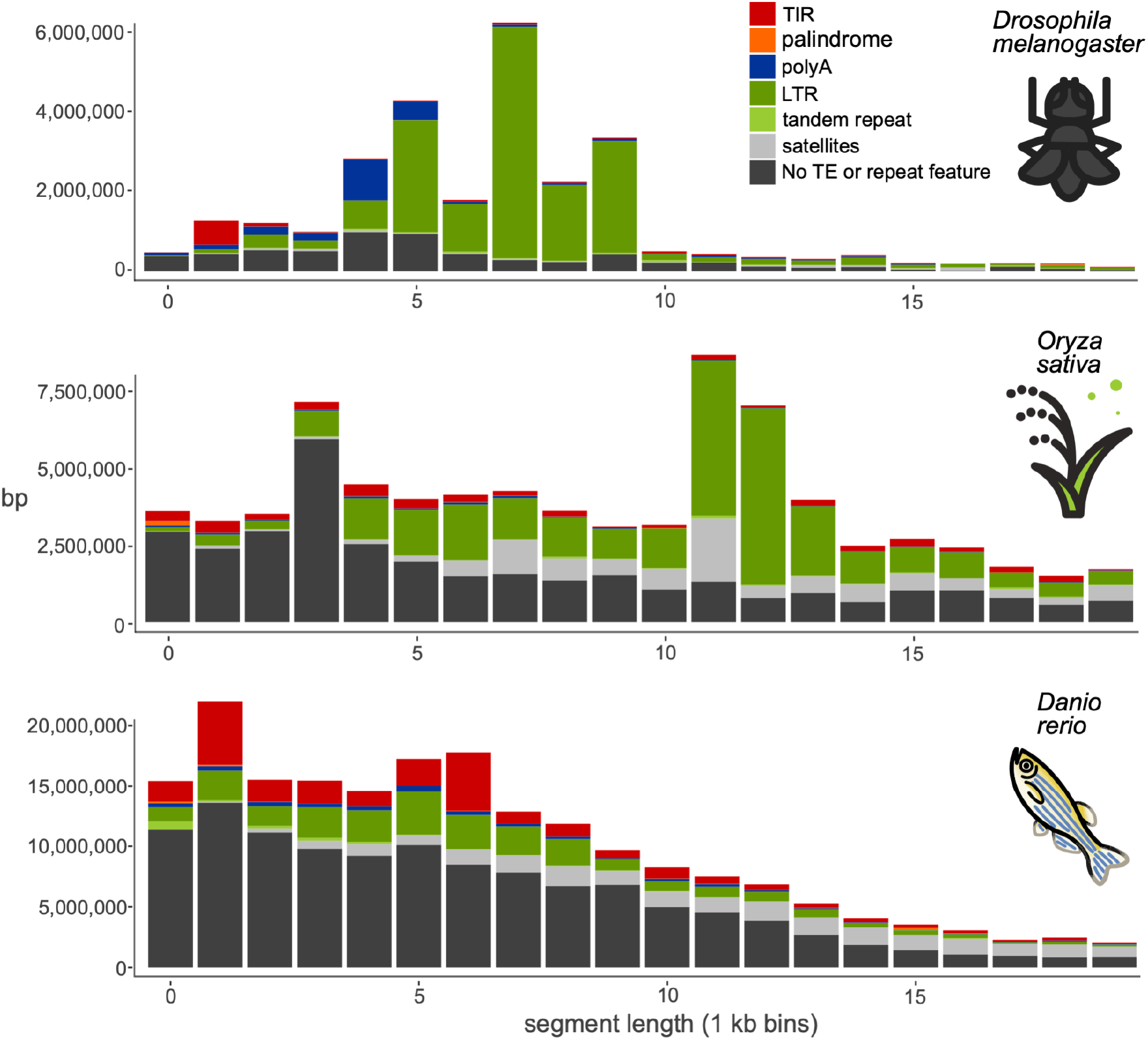
Structural feature found in polymorphic segments of three pangenomes, by segment length (250 - 20,000 bases). Total number of base pairs in insertions on a pangenome in different size ranges, grouped by structural features associated with TEs or other repeats. TIR: terminal inverted repeats. Palindrome: TIRs that occupy more than 90% of the sequence. polyA: A/T homopolymer at least 10 bases long, allowing for 1 mismatch every 8 bases. Tandem repeat: The sequence is composed of a smaller sequence repeated 2 or 3 times. Satellites: The sequence is composed of a motif repeated more than 3 times.

### Drosophila melanogaster

We used seven genomes A1 to A7 of *Drosophila melanogaster* from the DrosOmics project (Coronado-Zamora et al., 2023) (Figure 1a) and built a pangenome composed of one connected graph for each of the five main Muller elements (chromosome arms 2L, 2R, 3L, 3R and X). This produced 5 GFA files between 100 MB and 144 MB of size composed of a total of 5,564,238 segments. Running pantera on them resulted in a library of 141 elements that we classified using RepeatClassifier 2.0.4 (Flynn et al., 2020) (Figure 1c). Next we ran RepeatModeler 2.0.4 (Flynn et al., 2020) with default parameters on one of the genomes (A1) and compared the results (N=361) with pantera and the reference library.

### Oryza sativa

For rice we constructed the pangenome from two genome sequences: the reference genome GCF_001433935.1 from the Japonica group (Kawahara et al., 2013) and an Indica group genome GCA_001623345.3 (Zhang et al., 2016). The final graph was composed of 7,801,181 segments. The resulting library obtained with pantera had 525 elements of which 267 (51%) are classified by RepeatClassifier as Unknown. We compared this library to the manually curated TE annotation in Rice (v6.9.5) (Ou et al., 2019), with 2,431 elements. In this case we compared the results of pantera to the uncurated library obtained with REPET (Quesneville et al., 2003) (Quesneville et al., 2005) (Flutre et al., 2011) downloaded directly from REPETDB (Amselem et al., 2019) composed of 2,479 families.

### Danio rerio

For zebrafish we used the reference genome danRer11 (GCF_000002035.6) (Howe et al., 2013) and compared it to fDanRer4.1 (GCA_944039275.1), one of the recent assemblies generated by the Wellcome Sanger Institute Tree of Life programme. The final graph was composed of 32,943,885 segments. The library obtained from it using pantera returned 913 putative TE families with 29 (3%) of them being classified as unknown. We compared it to the 1,740 curated TE families included in Dfam (Storer et al., 2021), and to the results obtained with RepeatModeler2 (3,728 families).

### Benchmark results

To compare the results we looked at different values (Figure 3): a) how many sequences from the reference library were matched in at least 90% of the sequence by a sequence of the other tools; b) what fraction of the sequences obtained for each type were complete; c) the total percentage of the genome masked by the resulting libraries.

**Figure 3:**
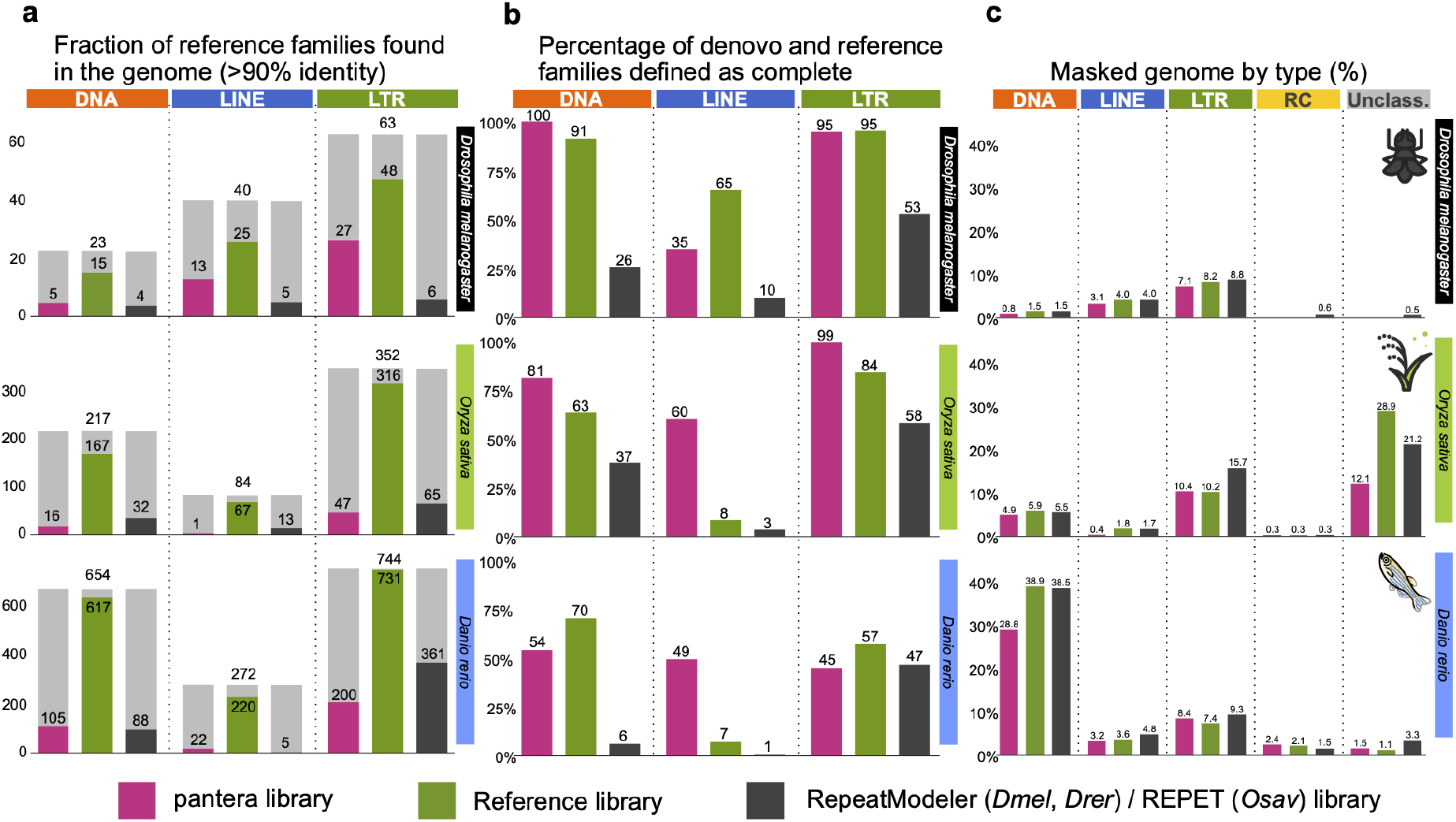
Comparing different TE libraries in *Drosophila melanogaster, Oryza sativa and Danio rerio*. **a,** Total number of families of the reference library (grey bars) and for each library how many of them have a match covering >90% identity and length in the selected genome (A1 for *Drosophila*, the reference for rice and zebrafish). Note that not all reference library families were found in the specific genome used (some are only found in other genomes from the species). **b,** Degree of TE completeness as percentages of the total number of segments for each tool, type and species. We define “complete” as: to have terminal inverted repeat (TIR) sequences for DNA elements (except Cryptons) and LTR DIRS elements; to have long terminal repeat (LTR) sequences for LTR elements; to have a candidate ORF at least 1300 aminoacids long, and a polyA tail for LINE elements. LINE1 elements in addition must contain an ORF1 candidate at least 700 aminoacids long. **c**, Percentage of the genome masked by RepeatMasker using each of the libraries by type of TE family.

Pantera found more near-full length (>90%) members of the reference libraries than RepeatModeler (fruit fly and zebrafish) or REPET (rice) except for LTR elements for rice and zebrafish. In rice this was primarily due to different criteria on divergence while defining a family, as was confirmed by the similar percentage of genome masked in both cases (pantera 10.4%, reference 10.2%). In the case of zebrafish both pantera and RepeatModeler libraries have an excess of incomplete elements, probably due to the relatively low copy number of full length LTR elements.

In general pantera families are more complete as defined in the previous section, even than the reference library families (Figure 3b), with the exceptions being DNA and LTR elements for zebrafish. Length distributions of all families generated can be compared in Figure 4 and Supp. Figure 1. We interpret the typically longer mean size and lower variance of the distributions of lengths by superfamilies for pantera as further evidence that the consensus sequences it produces tend to belong to full elements. As an example, of 48 CMC-EnSpm families identified by pantera, only 5 lack the expected TIR elements, compared to 29 families missing the TIR element out of 70 in the REPET results. This is even true for LINE elements, for which is particularly hard to produce a full length consensus because most copies are incomplete. Another point to take into account is that the results can also be biassed by the cut point selected to define the minimum size of an element to be included in the library. Mobile elements associated with TE activity usually start over the 100 bases mark, with SINEs or solo LTRs. If instead we want to focus on autonomous TEs, in our experience a minimum size of 700 to 800 bases is low enough. As pantera uses the information from several genomes, it is possible that a family found in the pangenome is not actually present in one of the genomes. This happens for example with the full LINE/CR1 element in fruit fly, that is in the curated and pantera libraries, but not present in the genome (A1) used by RepeatModeler and as template for the results.

**Figure 4:**
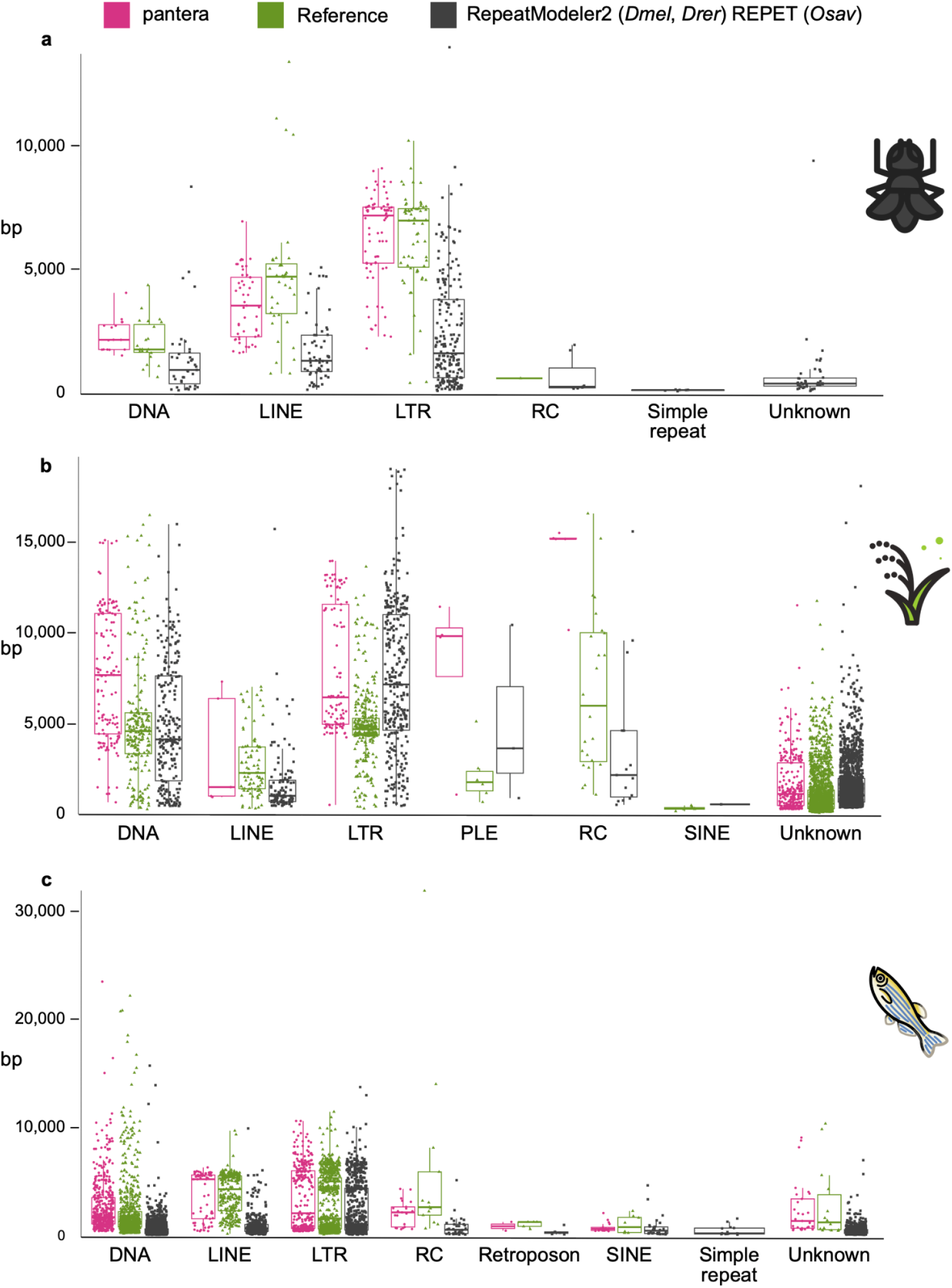
Length distributions of the different libraries by TE order. Length distributions of the consensus sequences by order in which they have been classified. RC stands for rolling circle (Helitrons). **a***, Drosophila melanogaster.* pantera (N=141), Drosophila Transposon Canonical Sequences 10.2 (N=127), RepeatModeler (N=361). **b***, Oryza sativa.* pantera (N=525), rice6.9.5 (N=2431), REPET (N=2471) **c***, Danio rerio.* pantera (N=913), Dfam curated (N=1740), RepeatModeler (N=3728).

The results of masking the genomes with the libraries generally show a comparable though slightly lower coverage percentage by pantera (Figure 3c). This is expected as pantera will not build consensus sequences from very old and fragmented TE insertions, that can represent a sizable percentage of the genome, and instead will identify more recent elements, which are closer to the putative active sequence of the TE, but which may have fewer copies in the genome. As an example, in *Danio rerio* pantera identified one large CMC-EnSpm element that has three full copies in the genome. It shows the two full proteins associated with these elements, and has a 13 basepair TIR (CACTCAAAAAAAT) (Supp. Fig. 2). This and other large CMC (CACTA) elements were not reported by RepeatModeler. The same happened with other large DNA elements classified as Zisupton (Supp. Fig. 3).

RepeatMasker landscape plots for all libraries are shown in Supp. Figures 4,5 and 6. Differences between methods are observed due to different clustering approaches. In general pantera provides greater resolution at low Kimura divergences, presumably due to its tight initial clustering step. We note that for *Danio rerio*, the landscapes for the curated library and the RepeatModeler library are very similar, suggesting perhaps that a RepeatModeler library was the basis for the curated library and that little reclustering was performed.

We compared the time employed by all workflows (Table 1) except for the REPET library for rice for which we used a library previously generated. For the pantera workflow we added the time employed by the creation of the pangenome (pggb) extraction of the library (pantera) and the classification of the sequences (RepeatClassifier). The results for RepeatModeler include also the time employed by RepeatClassifier. With *Drosophila melanogaster* pantera was 3.5x times faster, even though the pangenome was composed of 7 genomes. In the case of *Danio rerio* the pantera workflow was 6x times faster than RepeatModeler. It is worth noting that by default RepeatModeler limits the genome sampled to 400 MB. This limit can be increased to sample the full genome and avoid missing low copy elements, but it comes at a larger cost in execution time. The results presented used the default configuration.

**Table 1:** Run times for library creation workflows. Tests were performed on the University of Cambridge HPC facility using Dell PowerEdge XE8545 servers using 3rd generation AMD EPYC 64-core CPUs.

|  |  | *Time* | *cores* |
| --- | --- | --- | --- |
| *D. melanogaster* | Pangenome | 1h:32m | 8 |
|  | pantera | 10m | 8 |
|  | RepeatClassifier | 17m | 1 |
|  | Total pantera workflow | 1h:59m |  |
|  | RepeatModeler2 | 7h:28m | 16 |
| *O. sativa* | Pangenome | 2h:57m | 16 |
|  | pantera | 1h:05m | 16 |
|  | RepeatClassifier | 47m | 1 |
|  | Total pantera workflow | 4h:49m |  |
| *D. rerio* | Pangenome | 6h:24m | 24 |
|  | pantera | 1h:48m | 24 |
|  | RepeatClassifier | 1h:04m | 1 |
|  | Total pantera workflow | 9h:16m |  |
|  | RepeatModeler2 | 61h:32m | 32 |

### Results using both haplotypes of the same sample: Trachurus trachurus and Aquila chrysaetos

As an example of the application of pantera to a newly sequenced species without a reference we selected two species from the Sanger Institute, the Atlantic horse mackerel *Trachurus trachurus* and the golden eagle *Aquila chrysaetos*. For *T. trachurus* we used its primary (GCA_905171665) and alternate (GCA_905171655) haplotype assemblies (Genner, 2022) to extract a new TE library for the species using the pantera pipeline (1301 families). Then we compared the results with the Ensembl annotation for the species, obtained with RepeatModeler, without further manual curation (3718 families) (Figure 5). The results of masking the genome with both libraries are similar, but in the case of pantera more than double the elements in all three main divisions (DNA, LINE, LTR) appear to represent the full sequence of the TE. For example, pantera was able to found a new ERV element, 12,371 bases long, with 707 bases LTRs and two ORFs of 1402 and 1032 amino acids. There is just one full copy in the main haplotype and this is a case in which pantera can benefit from using the information present in both haplotypes (Figure 5d,e). Furthermore, the largest family of CMC-EnSpm elements found in the genome has no full copies in the primary haplotype but is only present in the alternate haplotype with 6 full copies (Figure 5f,g,h).

**Figure 5:**
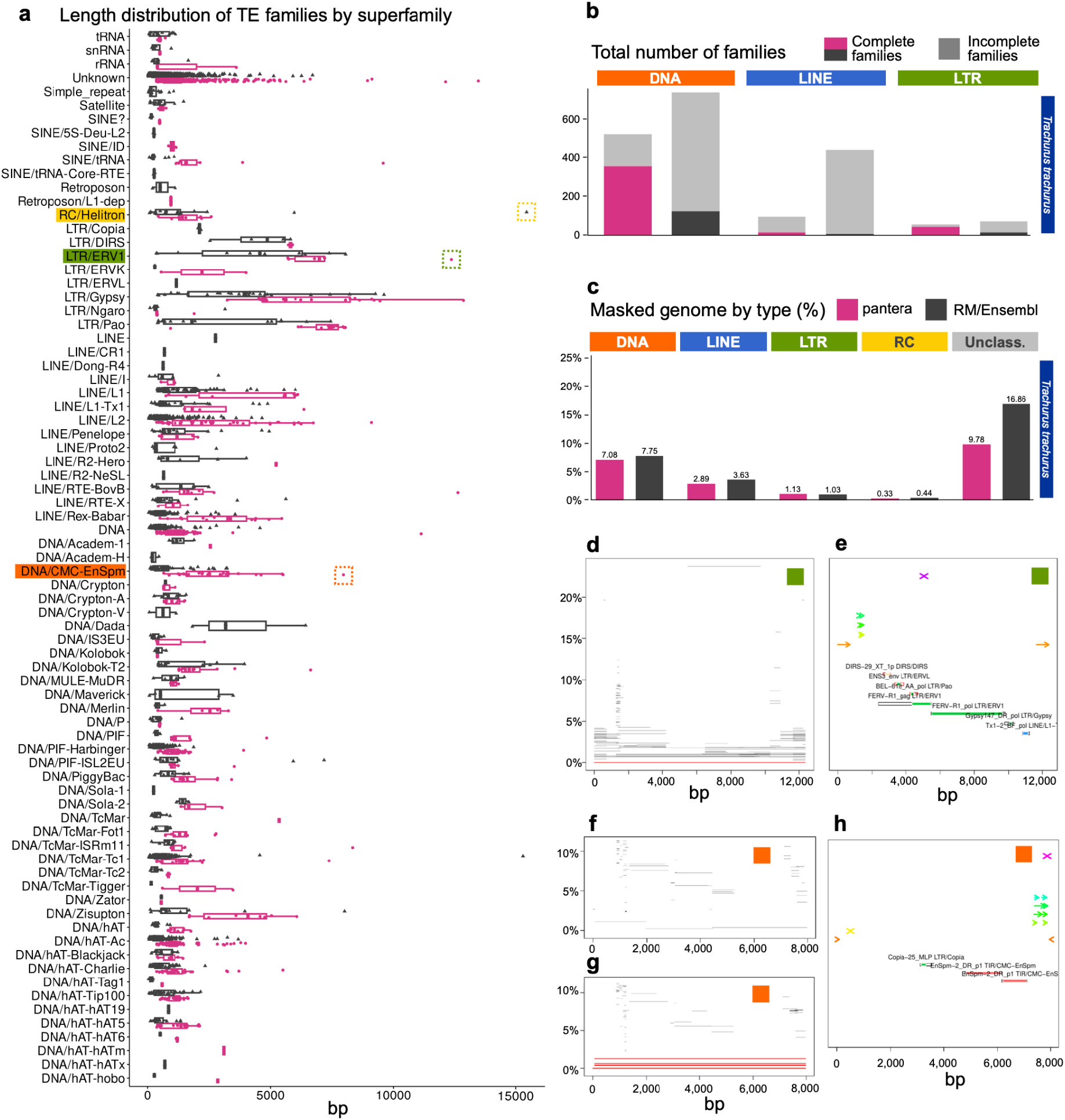
Results with *Trachurus trachurus*, from a pangenome composed of the primary and alternate assemblies from the same sample. **a**, Length distributions of the consensus sequences by superfamily in which they have been classified. RC stands for rolling circle (Helitrons) pantera (N = 1301), RepeatModeler (N=3718). **b,** Total number of families of the resulting libraries, and their degree of completeness as defined in the following way: We define “complete” as in Figure 2. **c,** Percentage of the genome masked by RepeatMasker using each of the libraries. **d,e,** Example of large ERV elements correctly identified by pantera. Just one copy is present in the principal haplotype. **h,** CACTA element found the alternate haplotype (**g**) but not in the principal haplotype (**f**).

We repeated the same procedure with the primary haplotype (GCA_900496995.4) and alternate (GCA_902153765.2) of the golden eagle (Mead et al., 2021). In this case pantera did not find any of the DNA type content found by RepeatModeler, as that appears to be due to old insertions which are no longer polymorphic. Instead, it was able to correctly find several large ERVs that are still polymorphic and might be relatively recent insertions, which were missed by RepeatModeler. In particular one of them includes an extra protein in addition to the putative ERV proteins that we found to be present also in the genomes of other Accipitriformes but not in more divergent species, which suggests that it could be an ERV specific to this order (Supp. Fig. 7).

### Results comparing closely related species: Astatotilapia calliptera and Maylandia zebra

The polymorphism-first approach can also be applied to comparisons between genomes of closely related species, and we have found that in some cases this allows us to have a better understanding of their TE content. As an example, we created a pangenome from the genomes of two closely related cichlid fishes from Lake Malawi, *Astatotilapia calliptera* (GCA_900246225.5) and *Maylandia zebra* (GCA_000238955.5), and used pantera to generate 250 candidate TE families. In the resulting library we found three different complete families of Maverick elements for which previously only fragmented components had been reported. One of them, Maverick-3_AstCal, has just one full copy in the *Astatotilapia calliptera* reference genome (Fig. 6a,b), but a search for polymorphic insertions in more than 600 samples with short read data using MeGANE (Kojima et al., 2022) confirmed that all of them have tens of polymorphic insertions of that family, highlighting the relevance of having the most complete possible consensus sequence to perform further downstream analysis accurately (Fig. 6c,d). Pantera also found a previously identified element, named piggybac-5, formed by the fusion of two segments of the same piggybac-like element in opposite senses (Fig. 6e,f,g). This has lost the transposase, but the intact TIRs suggest it is still being mobilized as a nonautonomous element, and indeed there are 51 full length copies in the *Astatotilapia calliptera* reference genome. Pantera also obtained a consensus for an intact piggybac TE (TE-243928) (Fig. 6f) which has only six full copies in the *Astatotilapia calliptera* genome, each containing a complete piggybac transposase of 256 aa. The target site duplications of TE-243928 and piggybac-5 are identical, and the terminal region of the TIR of piggybac-5 is the same as the TIR of TE-243928, but the piggybac-5 TIR is substantially extended internally by material which is only found in single copy in TE-243928 (Fig. 6h). We suggest that piggybac-5 may have been formed by overlapping chromosomal inversion events from TE-243928 or a closely related element.

**Figure 6:**
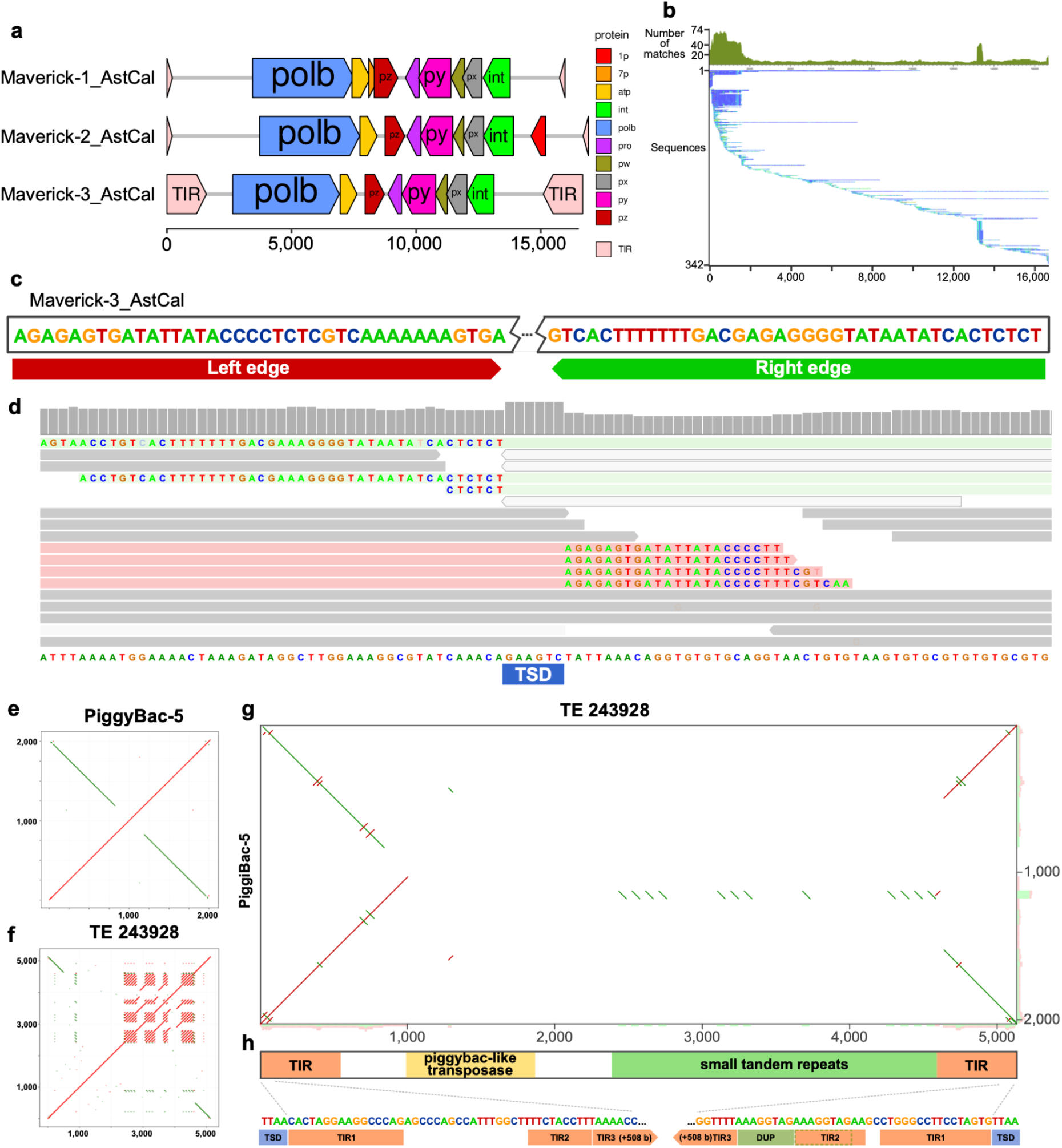
Selected TE families found in *Astatotilapia calliptera*. **a**, Structure elements identified in three Maverick families found with pantera in a pangenome built with *Astatotilapia calliptera* and *Maylandia zebra*. All families include TIR elements, and separate ORF components for DNA polymerase b, integrase, ATPase and a double jelly roll capsid protein (py) among others. **b**, Seed Alignment Coverage and Whisker Plot for Maverick-3_AstCal. Only one full copy is present in the reference genome, which makes it basically impossible to find it based only on repetitiveness. **c**, Detail of the edge sequences for both TIR elements. **d**, The accurate definition of the edges of the TE element allows us later to use other tools like MeGANE to identify polymorphic insertions using short reads, bases on mapping of discordant reads to the TE sequence but also matching the soft clip reads of on the insertion to the edges of the putative TE element. In this case we observe the signal for an heterozygous polymorphic insertion, which has created an 8 bases target segment duplication (TSD). **e**, Structure of a new TE composed of the fusion of two identical piggybac elements in opposite sense. **f**, Hits in the genome. The black divergent lines show matches to previous insertions of the single piggybac element. The red complete ones prove that the new element is creating new copies. **g**, Self dotplot showing the structure of the element. **f** and **g** generated with TE-aid (https://github.com/clemgoub/TE-Aid).

## Discussion

The pros and cons of pantera are related to the “polymorphism first” approach. A benefit is that the consensus sequences obtained are often closer to the curated sequences than with standard “repeat first” approaches, when those sequences belong to recent elements in which the structural features are still intact. This is reflected also in obtaining a smaller percentage of sequences that could not be assigned to any category (Sup Tables 2-4). This is the result of the initial selection of segments (polymorphic and highly similar), that are more likely to be the product of recent transposition events and so reflect full TE elements. Unlike methods starting from a seed that is extended until the final consensus is found, our method is less constrained by size, as the size of the segments found is a direct result of the pangenome construction, and many of the segments originally selected are expected to represent the full sequence of the TE. Another benefit is that it does not need a large number of repetitive elements to identify the putative sequences, relying instead on the presence of at least two polymorphic copies in the pangenome.

We show several examples where Pantera identified TE families missed by other automated methods (Figures 5d-h, 6b). In particular, because it initially clusters full length sequences with high stringency, it can distinguish related separately transposing sequences that share common sections, illustrating how novel elements can arise from fragments of previous ones (Figure 6e-h).

The tendency to generate full length consensus sequences is valuable for downstream tools which require an accurate library to genotype TEs polymorphisms [41], particularly in species that contain hundreds or thousands of TE families, making it a daunting prospect to manually curate them. These tools to genotype TEs based on short reads usually are based on the information obtained from discordant read mapping to TEs and on the unmapped content of split reads. In both cases it is necessary to use the full TE consensus to accurately assign a polymorphism to that family. (Figure 6-c,d)

A limitation of the applicability of pantera is that, since it is a comparative method, it requires more than one genome sequence. Furthermore, elements which are not polymorphic between the genomes will not be identified. It might be expected that this would restrict pantera to only finding very recently transposed sequences, but as we saw in our evaluation it is surprisingly successful at generating a library that masks a similar fraction of the genome as more traditional approaches. It seems that for old insertions this is typically achieved not by including a consensus for the original transposon in the library, but rather by masking with a more recently active descendant (or other close relative). We note that such a relative is normally no more divergent from the original TE than the actual insertions themselves - both will have drifted by mutation since their shared common ancestor. However, we recognise that ideally for old insertions it would be preferable to use a consensus that more accurately reflects the original active transposon sequence.

For this and other reasons, we do not claim that pantera by itself will replace existing approaches. Instead we suggest that it has complementary properties to them and will provide a valuable addition to composite TE annotation approaches such as EarlGrey alongside repeat-first methods. Because pantera tends to generate full length consensus sequences more frequently than tools that start from a repeat-first approach, we suggest that it might be used first, then the genome be masked for sequences found by pantera, then a method such as RepeatModeler or REPET be used on the masked genome.

## Conclusions

We present a novel approach to the identification of TEs based on insertion polymorphism, together with a practical software implementation, pantera. The results of this approach are complementary to those of previous automated transposon family discovery tools and can be used to reduce the curation required to build a high quality transposon library. To make libraries for a new species it relies on there being multiple assemblies of a genome from different haplotypes, but these are now standard from modern long-read genome assemblies as generated for example by the Vertebrate Genomes Project [42], the Darwin Tree of Life project [15] or other Earth Biogenome Project sequencing projects. As more species have their genomes sequenced it will be increasingly possible to apply pantera between closely related species.

## Data availability

The code for pantera and all the libraries generated by it discussed in this paper are available from https://github.com/piosierra/pantera. The *Drosophila melanogaster* genome sequences that were used can be downloaded from https://www.biologiaevolutiva.org/gonzalez_lab/drosomics/DATA/. Accession numbers for *O. sativa* and *D. rerio* genome sequences are given in the text.

## Acknowledgments

The authors want to acknowledge the support of Simon Orozco (Institut de Biologia Evolutiva - CSIC UPF) for his feedback on TE libraries benchmarking methods, Johann Confais (INRAE) for providing the REPET library for *Oryza sativa* and Jessica Storer (Dfam - University of Connecticut) for her insights on TE curation.

*Drosophila melanogaster*, *Danio rerio* and *Oryza sativa* icons were obtained from TogoTV (© 2016 DBCLS TogoTV, CC-BY-4.0 https://creativecommons.org/licenses/by/4.0/).

## Author contributions

P.S. and R.D. conceived the project and wrote the paper, P.S. implemented pantera, carried out the experiments, and analysed the results with the assistance of R.D.

## List of abbreviations

GFA: Graphical fragmented assembly format
LTR: Long terminal repeat
SV: Structural variant
TE: Transposable element
TIR: Terminal inverted repeat
TSD: Target site duplication

## Supplementary Figures

**Supplementary Figure 1:**
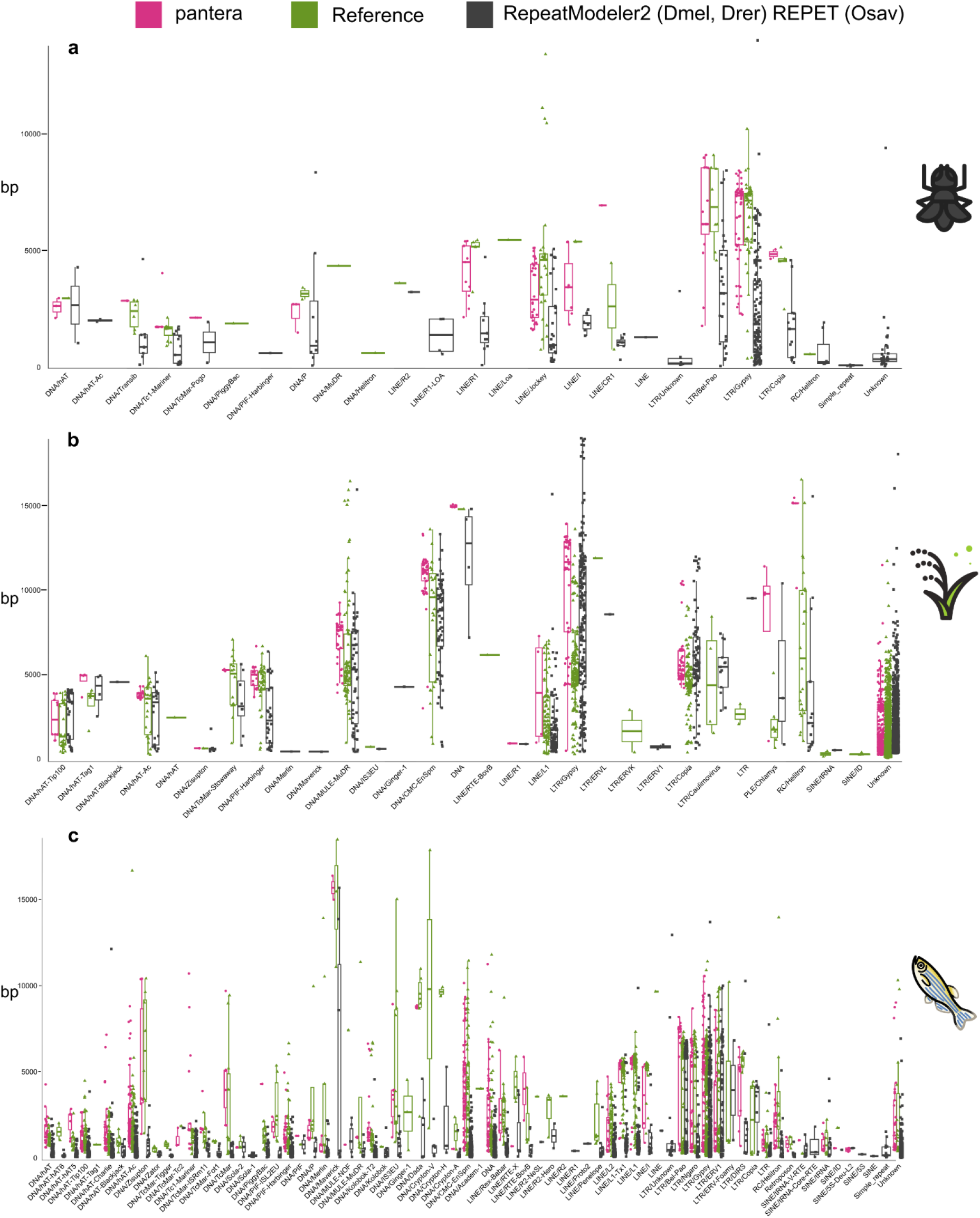
Length distributions of the different libraries by TE superfamily. Length distributions of the consensus sequences by superfamily in which they have been classified. RC stands for rolling circle (Helitrons). **a***, Drosophila melanogaster.* pantera (N=141), Drosophila Transposon Canonical Sequences 10.2 (N=127), RepeatModeler (N=361). **b***, Oryza sativa.* pantera (N=525), rice6.9.5 (N=2431), REPET (N=2471) **c***, Danio rerio.* pantera (N=913), Dfam curated (N=1740), RepeatModeler (N=3728).

**Supplementary Figure 2:**
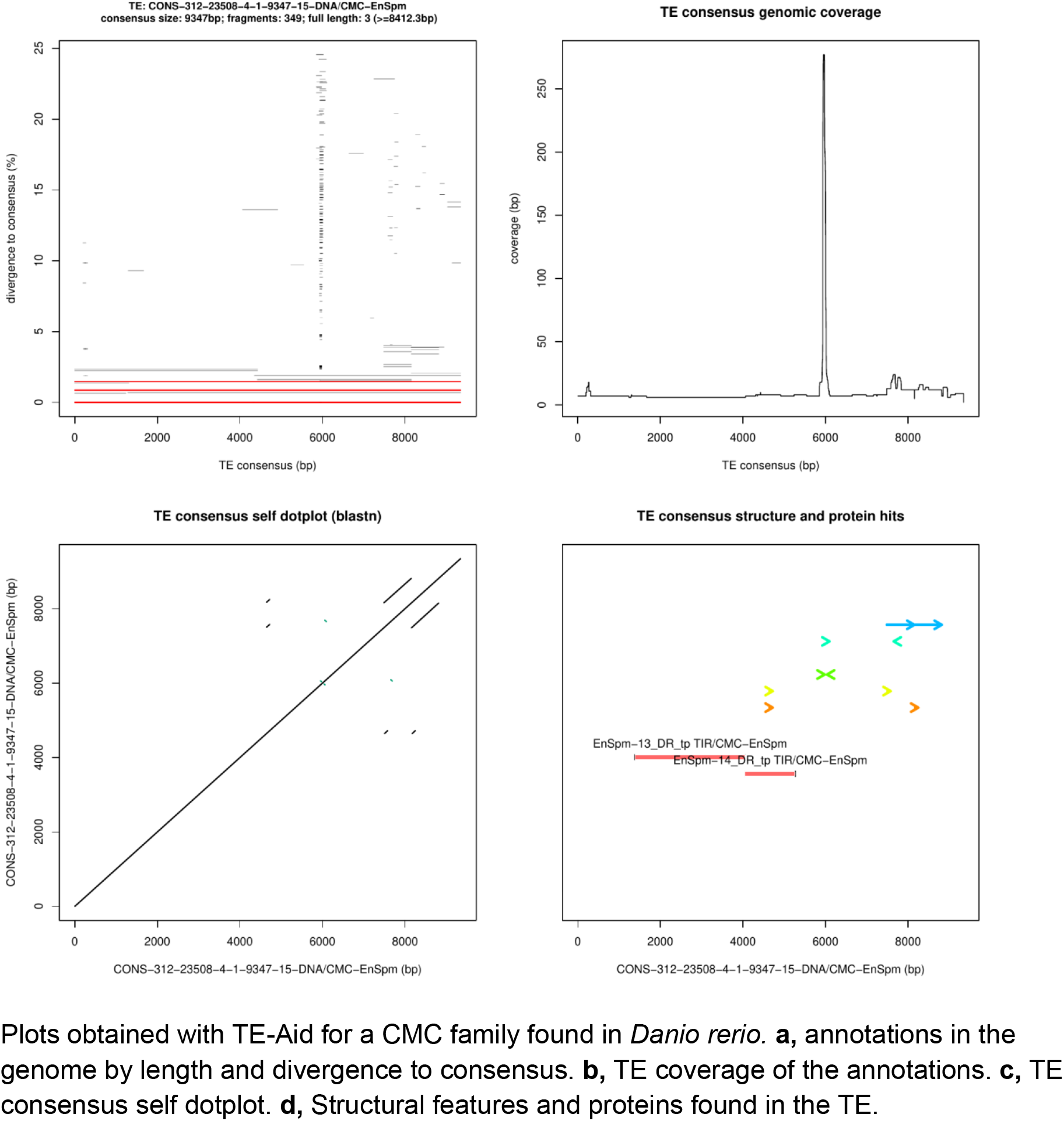
CMC-EnSpm family from *Danio rerio*. Plots obtained with TE-Aid for a CMC family found in *Danio rerio.* **a,** annotations in the genome by length and divergence to consensus. **b,** TE coverage of the annotations. **c,** TE consensus self dotplot. **d,** Structural features and proteins found in the TE.

**Supplementary Figure 3:**
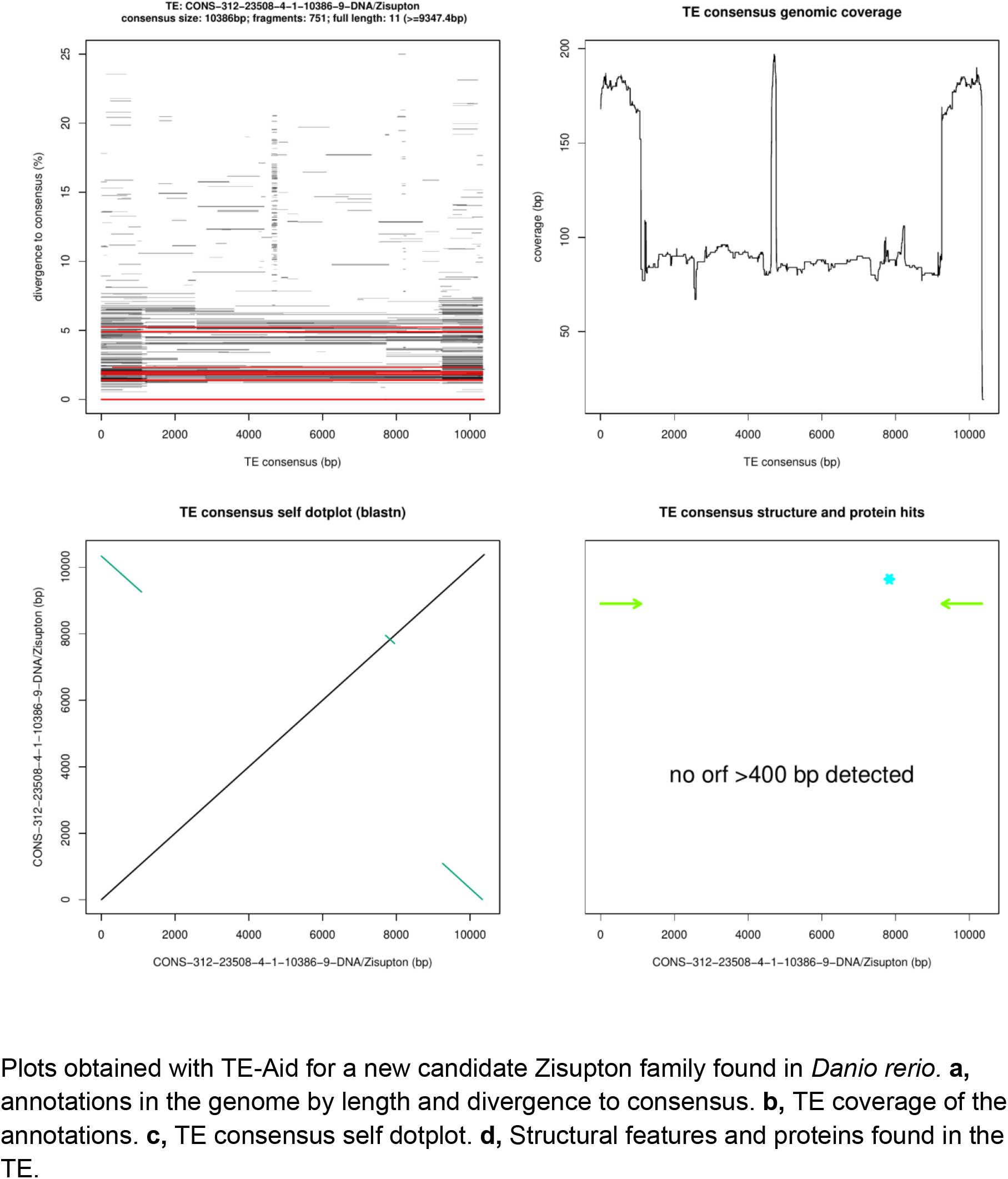
Zisupton candidate family from *Danio rerio*. Plots obtained with TE-Aid for a new candidate Zisupton family found in *Danio rerio.* **a,** annotations in the genome by length and divergence to consensus. **b,** TE coverage of the annotations. **c,** TE consensus self dotplot. **d,** Structural features and proteins found in the TE.

**￼￼￼Supplementary Figure 4:**
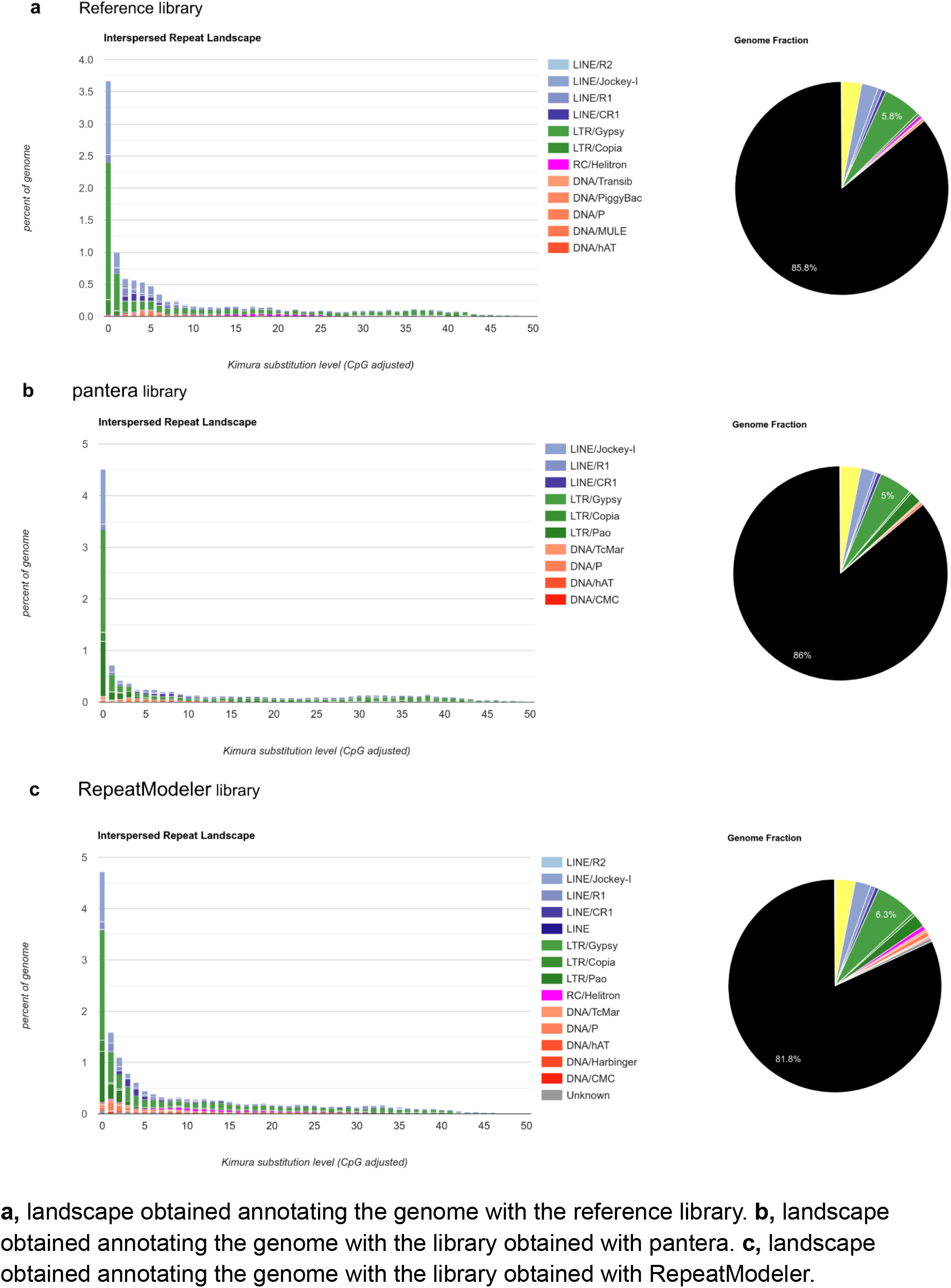
RepeatMasker generated TE landscapes for *Drosophila melanogaster.* **a,** landscape obtained annotating the genome with the reference library. **b,** landscape obtained annotating the genome with the library obtained with pantera. **c,** landscape obtained annotating the genome with the library obtained with RepeatModeler.

**Supplementary Figure 5:**
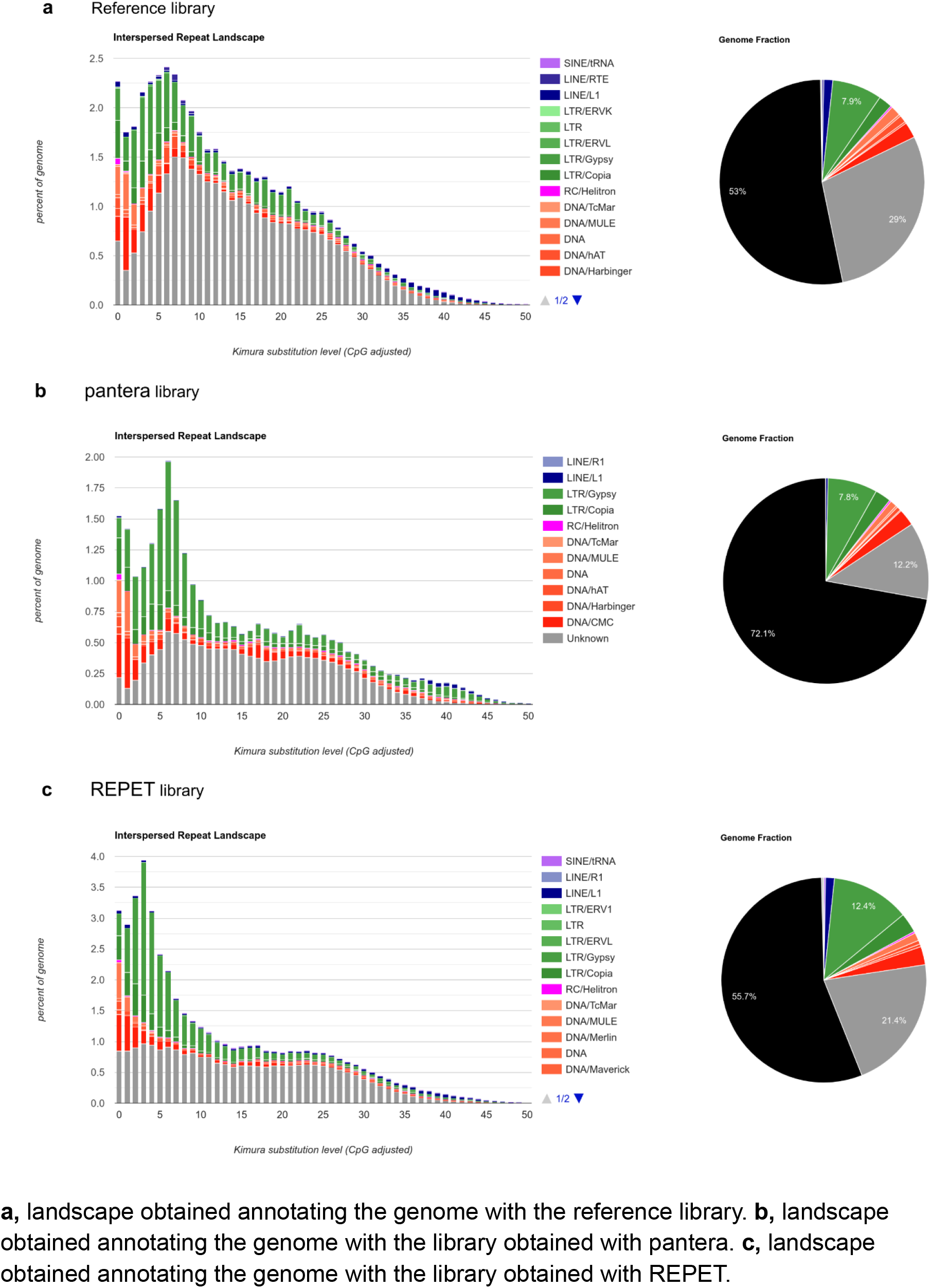
RepeatMasker generated TE landscapes for *Oryza sativa*. **a,** landscape obtained annotating the genome with the reference library. **b,** landscape obtained annotating the genome with the library obtained with pantera. **c,** landscape obtained annotating the genome with the library obtained with REPET.

**Supplementary Figure 6:**
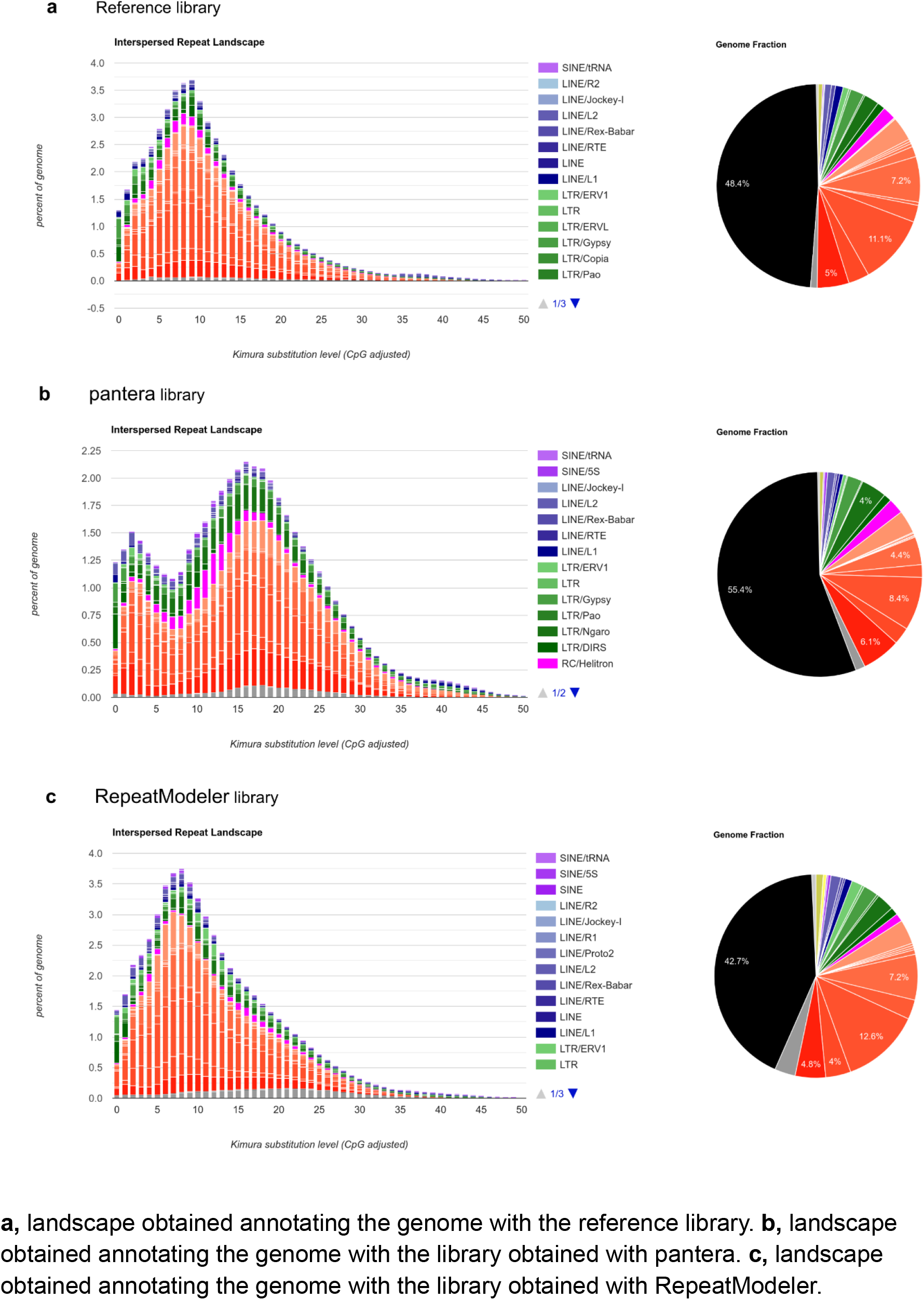
RepeatMasker generated TE landscapes for *Danio rerio*. **a,** landscape obtained annotating the genome with the reference library. **b,** landscape obtained annotating the genome with the library obtained with pantera. **c,** landscape obtained annotating the genome with the library obtained with RepeatModeler.

**Supplementary Figure 7:**
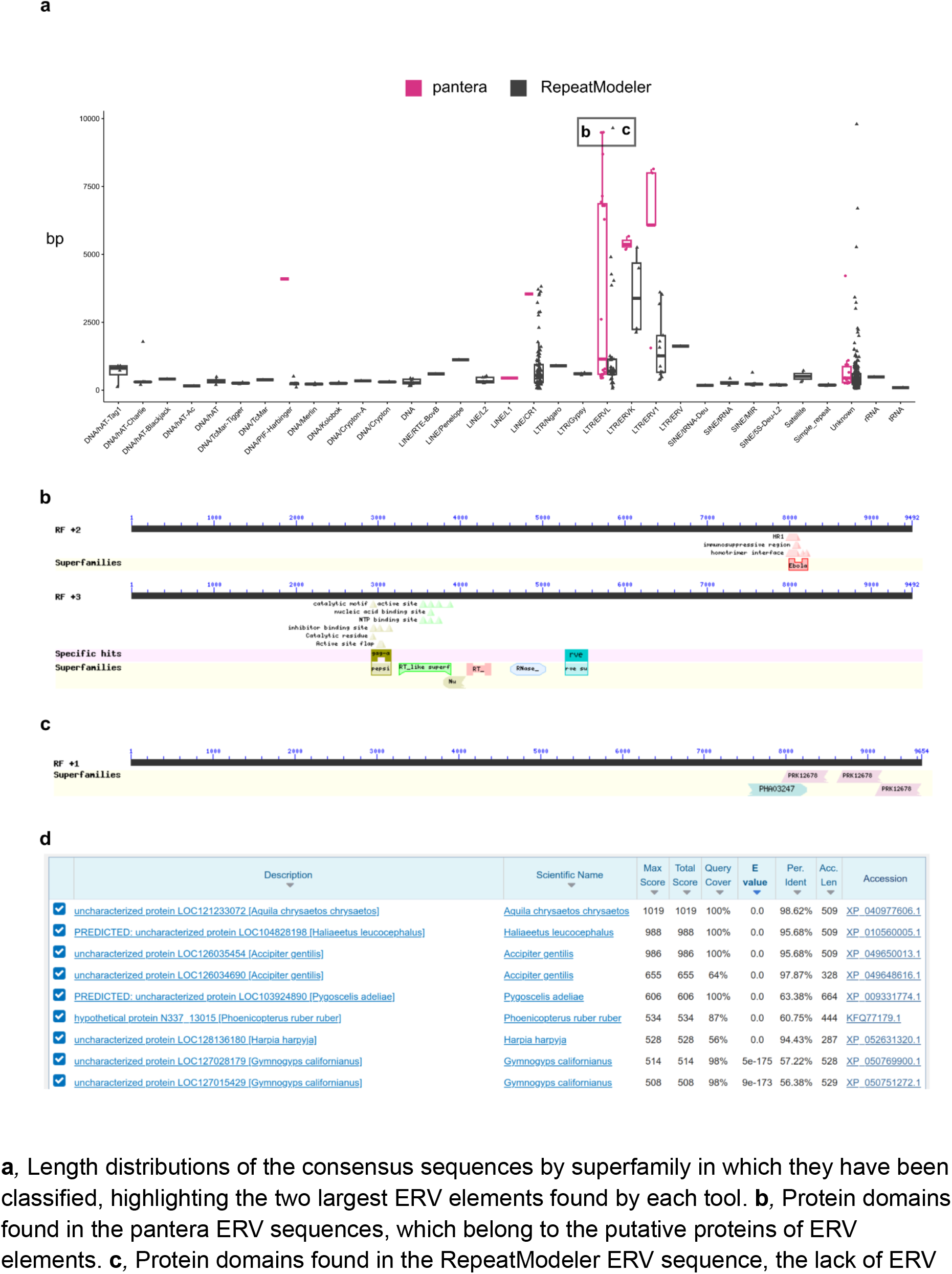
Example of correct and incorrectly annotated TEs (large ERVs) in the TE libraries for *Aquila cryasetos*. **a**, Length distributions of the consensus sequences by superfamily in which they have been classified, highlighting the two largest ERV elements found by each tool. **b**, Protein domains found in the pantera ERV sequences, which belong to the putative proteins of ERV elements. **c**, Protein domains found in the RepeatModeler ERV sequence, the lack of ERV related proteins suggest it is either an artefact or the consensus has not been properly resolved. **d**, The same ERV protein can be found in other species of the same clade.

**Supplementary Figure 8:**
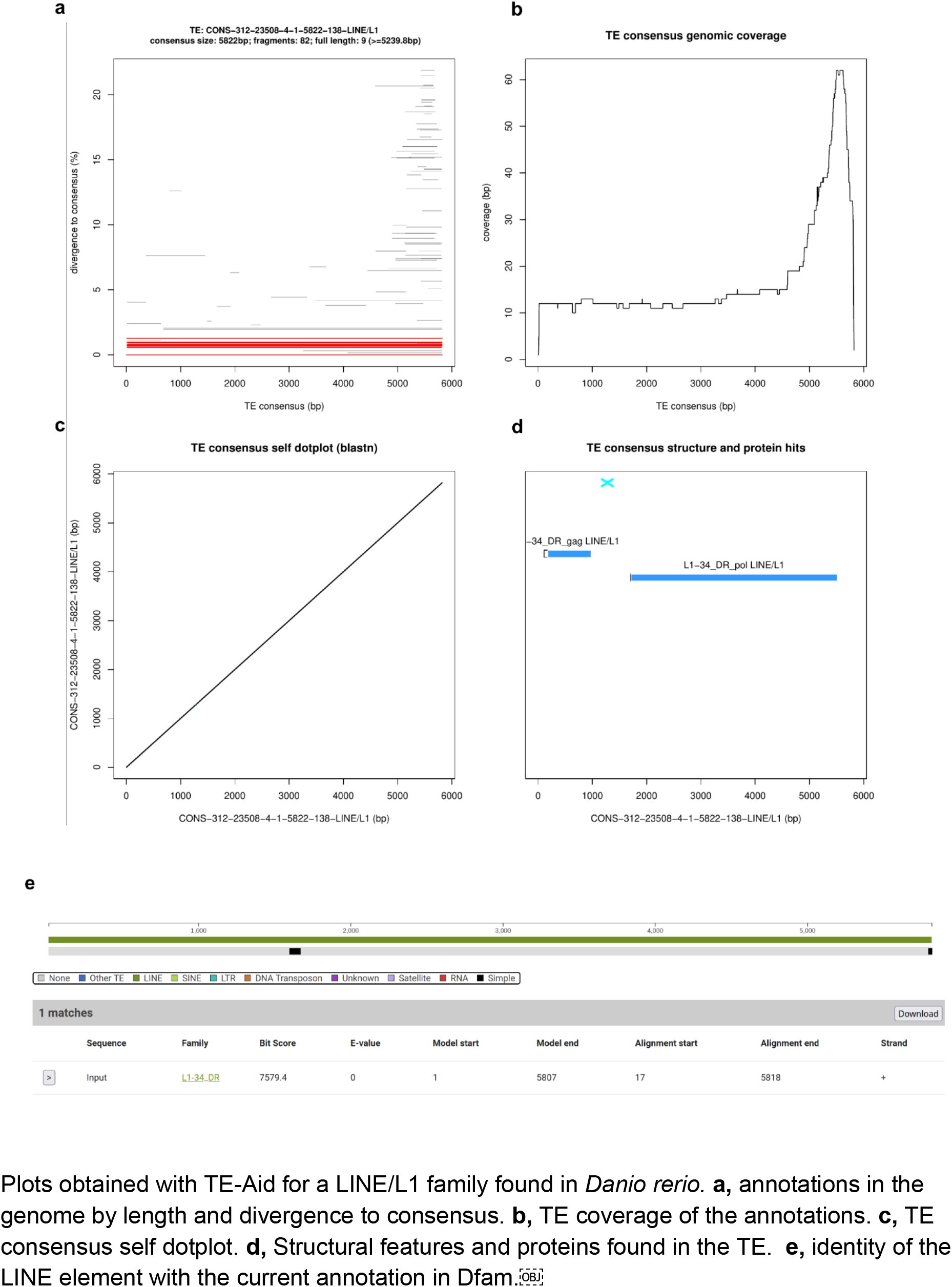
LINE/L1 family from *Danio rerio*. Plots obtained with TE-Aid for a LINE/L1 family found in *Danio rerio.* **a,** annotations in the genome by length and divergence to consensus. **b,** TE coverage of the annotations. **c,** TE consensus self dotplot. **d,** Structural features and proteins found in the TE. **e,** identity of the LINE element with the current annotation in Dfam.￼

## References

Amselem, J., Cornut, G., Choisne, N., Alaux, M., Alfama-Depauw, F., Jamilloux, V., et al. (2019). RepetDB: a unified resource for transposable element references. Mob. DNA 10, 1–8. doi: 10.1186/s13100-019-0150-y

Baril, T., Imrie, R. M., and Hayward, A. (2022). Earl Grey: a fully automated user-friendly transposable element annotation and analysis pipeline. 2022.06.30.498289.doi: 10.1101/2022.06.30.498289

Berthelier, J., Casse, N., Daccord, N., Jamilloux, V., Saint-Jean, B., and Carrier, G. (2018). A transposable element annotation pipeline and expression analysis reveal potentially active elements in the microalga Tisochrysis lutea. BMC Genomics 19, 1–14. doi: 10.1186/s12864-018-4763-1

Burke, D., Chuong, E., Taylor, W., and Layer, R. M. (2023). TEPEAK : A novel method for identifying and characterizing polymorphic transposable elements in non-model species populations. 2023.10.13.562297.doi: 10.1101/2023.10.13.562297

Cheng, H., Concepcion, G. T., Feng, X., Zhang, H., and Li, H. (2021). Haplotype-resolved de novo assembly using phased assembly graphs with hifiasm. Nat. Methods 18, 170–175. doi: 10.1038/s41592-020-01056-5

Coronado-Zamora, M., Salces-Ortiz, J., and González, J. (2023). DrosOmics: A Browser to Explore -omics Variation Across High-Quality Reference Genomes From Natural Populations of Drosophila melanogaster. Mol. Biol. Evol. 40, msad075. doi: 10.1093/molbev/msad075

Elliott, T. A., Heitkam, T., Hubley, R., Quesneville, H., Suh, A., and Wheeler, T. J. (2021). TE Hub: A community-oriented space for sharing and connecting tools, data, resources, and methods for transposable element annotation. Mob. DNA 12, 1–5. doi: 10.1186/s13100-021-00244-0

Flutre, T., Duprat, E., Feuillet, C., and Quesneville, H. (2011). Considering Transposable Element Diversification in De Novo Annotation Approaches. PLOS ONE 6, e16526. doi: 10.1371/journal.pone.0016526

Flynn, J. M., Hubley, R., Goubert, C., Rosen, J., Clark, A. G., Feschotte, C., et al. (2020). RepeatModeler2 for automated genomic discovery of transposable element families. Proc. Natl. Acad. Sci. 117, 9451–9457. doi: 10.1073/pnas.1921046117

Garrison, E., Guarracino, A., Heumos, S., Villani, F., Bao, Z., Tattini, L., et al. (2023). Building pangenome graphs. 2023.04.05.535718.doi: 10.1101/2023.04.05.535718

Genereux, D. P., Serres, A., Armstrong, J., Johnson, J., Marinescu, V. D., Murén, E., et al. (2020). A comparative genomics multitool for scientific discovery and conservation. Nature 587, 240–245. doi: 10.1038/s41586-020-2876-6

Genner, M. J. (2022). The genome sequence of the Atlantic horse mackerel. Wellcome Open Res. Open Access Publ. Platf. doi: 10.12688/wellcomeopenres.17813.1

Goubert, C., Craig, R. J., Bilat, A. F., Peona, V., Vogan, A. A., and Protasio, A. V. (2022). A beginner’s guide to manual curation of transposable elements. Mob. DNA 13, 7. doi: 10.1186/s13100-021-00259-7

Groza, C., Bourque, G., and Goubert, C. (2023). “A Pangenome Approach to Detect and Genotype TE Insertion Polymorphisms,” in Transposable Elements: Methods and Protocols Methods in Molecular Biology., eds. M. R. Branco and A. de Mendoza Soler (New York, NY: Springer US), 85–94. doi: 10.1007/978-1-0716-2883-6_5

Hickey, G., Heller, D., Monlong, J., Sibbesen, J. A., Sirén, J., Eizenga, J., et al. (2020). Genotyping structural variants in pangenome graphs using the vg toolkit. Genome Biol. 21, 1–17. doi: 10.1186/s13059-020-1941-7

Howe, K., Clark, M. D., Torroja, C. F., Torrance, J., Berthelot, C., Muffato, M., et al. (2013). The zebrafish reference genome sequence and its relationship to the human genome. Nature 496, 498–503. doi: 10.1038/nature12111

Kawahara, Y., de la Bastide, M., Hamilton, J. P., Kanamori, H., McCombie, W. R., Ouyang, S., et al. (2013). Improvement of the Oryza sativa Nipponbare reference genome using next generation sequence and optical map data. Rice 6, 1–10. doi: 10.1186/1939-8433-6-4

Kohany, O., Gentles, A. J., Hankus, L., and Jurka, J. (2006). Annotation, submission and screening of repetitive elements in Repbase: RepbaseSubmitter and Censor. BMC Bioinformatics 7, 1–7. doi: 10.1186/1471-2105-7-474

Kojima, S., Koyama, S., Ka, M., Saito, Y., Parrish, E. H., Endo, M., et al. (2022). Mobile elements in human population-specific genome and phenotype divergence. 2022.03.25.485726.doi: 10.1101/2022.03.25.485726

Li, H. (2018). Minimap2: pairwise alignment for nucleotide sequences. Bioinformatics 34, 3094–3100. doi: 10.1093/bioinformatics/bty191

Li, H., and Durbin, R. (2023). Genome assembly in the telomere-to-telomere era. doi: 10.48550/arXiv.2308.07877

Li, H., Feng, X., and Chu, C. (2020). The design and construction of reference pangenome graphs with minigraph. Genome Biol. 21, 1–19. doi: 10.1186/s13059-020-02168-z

Liao, W.-W., Asri, M., Ebler, J., Doerr, D., Haukness, M., Hickey, G., et al. (2023). A draft human pangenome reference. Nature 617, 312–324. doi: 10.1038/s41586-023-05896-x

Marco-Sola, S., Eizenga, J. M., Guarracino, A., Paten, B., Garrison, E., and Moreto, M. (2022). Optimal gap-affine alignment in O(s) space. 2022.04.14.488380. doi: 10.1101/2022.04.14.488380

McDavid, A., Gu, Y., VonKaenel, E., and Wagner, A. (2023). CellaRepertorium: Data structures, clustering and testing for single cell immune receptor repertoires (scRNAseq RepSeq/AIRR-seq). doi: 10.18129/B9.bioc.CellaRepertorium

Mead, D., Ogden, R., Meredith, A., Peniche, G., Smith, M., Corton, C., et al. (2021). The genome sequence of the European golden eagle, Aquila chrysaetos chrysaetos Linnaeus 1758. Wellcome Open Res. 6, 112. doi: 10.12688/wellcomeopenres.16631.1

Novák, P., Neumann, P., and Macas, J. (2010). Graph-based clustering and characterization of repetitive sequences in next-generation sequencing data. BMC Bioinformatics 11, 378. doi: 10.1186/1471-2105-11-378

Orozco-Arias, S., Sierra, P., Durbin, R., and González, J. (2023). MCHelper automatically curates transposable element libraries across species. 2023.10.17.562682. doi: 10.1101/2023.10.17.562682

Ou, S., Collins, T., Qiu, Y., Seetharam, A. S., Menard, C. C., Manchanda, N., et al. (2022). Differences in activity and stability drive transposable element variation in tropical and temperate maize. 2022.10.09.511471. doi: 10.1101/2022.10.09.511471

Ou, S., Su, W., Liao, Y., Chougule, K., Agda, J. R. A., Hellinga, A. J., et al. (2019). Benchmarking transposable element annotation methods for creation of a streamlined, comprehensive pipeline. Genome Biol. 20, 1–18. doi: 10.1186/s13059-019-1905-y

Quesneville, H., Bergman, C. M., Andrieu, O., Autard, D., Nouaud, D., Ashburner, M., et al. (2005). Combined Evidence Annotation of Transposable Elements in Genome Sequences. PLOS Comput. Biol. 1, e22. doi: 10.1371/journal.pcbi.0010022

Quesneville, H., Nouaud, D., and Anxolabéhère, D. (2003). Detection of New Transposable Element Families in Drosophila melanogaster and Anopheles gambiae Genomes. J. Mol. Evol. 57, S50–S59. doi: 10.1007/s00239-003-0007-2

Rautiainen, M., Nurk, S., Walenz, B. P., Logsdon, G. A., Porubsky, D., Rhie, A., et al. (2023). Telomere-to-telomere assembly of diploid chromosomes with Verkko. Nat. Biotechnol., 1–9. doi: 10.1038/s41587-023-01662-6

Rhie, A., McCarthy, S. A., Fedrigo, O., Damas, J., Formenti, G., Koren, S., et al. (2021). Towards complete and error-free genome assemblies of all vertebrate species. Nature 592, 737–746. doi: 10.1038/s41586-021-03451-0

Rice, P., Longden, I., and Bleasby, A. (2000). EMBOSS: the European Molecular Biology Open Software Suite. Trends Genet. TIG 16, 276–277. doi: 10.1016/s0168-9525(00)02024-2

Riehl, K., Riccio, C., Miska, E. A., and Hemberg, M. (2022). TransposonUltimate: software for transposon classification, annotation and detection. *Nucleic Acids Res.*, gkac136. doi: 10.1093/nar/gkac136

Smit, Hubley, R & Green, P. (2013). RepeatMasker. Available at: http://www.repeatmasker.org/RepeatMasker/ (Accessed December 30, 2020).

Storer, J., Hubley, R., Rosen, J., Wheeler, T. J., and Smit, A. F. (2021). The Dfam community resource of transposable element families, sequence models, and genome annotations. Mob. DNA 12, 1–14. doi: 10.1186/s13100-020-00230-y

Storer, J. M., Hubley, R., Rosen, J., and Smit, A. F. A. (2022). Methodologies for the De Novo Discovery of Transposable Element Families. Genes 13, 709. doi: 10.3390/genes13040709

The Darwin Tree of Life Project Consortium (2022). Sequence locally, think globally: The Darwin Tree of Life Project. Proc. Natl. Acad. Sci. 119, e2115642118. doi: 10.1073/pnas.2115642118

Wells, J. N., and Feschotte, C. (2020). A Field Guide to Eukaryotic Transposable Elements. Annu. Rev. Genet. 54, 539–561. doi: 10.1146/annurev-genet-040620-022145

Zhang, J., Chen, L.-L., Sun, S., Kudrna, D., Copetti, D., Li, W., et al. (2016). Building two indica rice reference genomes with PacBio long-read and Illumina paired-end sequencing data. Sci. Data 3, 160076. doi: 10.1038/sdata.2016.76

